# Metabolic regulation in erythroid differentiation by systemic ketogenesis in fasted mice

**DOI:** 10.1101/2023.06.07.544102

**Authors:** Wenjuan Ma, Yuichiro Arima, Terumasa Umemoto, Tomomasa Yokomizo, Yuqing Xu, Kenichi Miharada, Yosuke Tanaka, Toshio Suda

## Abstract

Systemic ketogenesis affects murine erythroid differentiation under fasting condition, while less ketone body β OHB boosts fatty acid synthesis and mevalonate pathway along with decreased levels of histone acetylation, which are beneficial for erythroid differentiation and maturation undergoing stressed erythropoiesis.

**Abstract:** Erythroid terminal differentiation and maturation depends on enormous energy supply. During periods of fasting, ketone bodies from the liver are transported into circulation and utilized as crucial fuel for peripheral tissues. However, the effects of fasting or ketogenesis on erythroid behavior remain unknown. Here, we generated a mouse model with insufficient ketogenesis by conditionally knocking out the gene encoding the hepatocyte-specific ketogenic enzyme hydroxymethylglutary-CoA synthase 2 (*Hmgcs2* KO). Intriguingly, erythroid maturation was enhanced with boosted fatty acid synthesis in bone marrow of hepatic *Hmgcs2* KO mouse under fasting condition, suggesting that systemic ketogenesis has a profound effect on erythropoiesis. Moreover, we observed significantly activated fatty acids synthesis and mevalonate pathway along with reduced histone acetylation in immature erythrocytes under less systemic ketogenesis condition. Our findings revealed an innovative insight to erythroid differentiation, in which metabolic homeostasis and histone acetylation mediated by ketone bodies are essential factors in adaptation towards nutrient deprivation and stressed erythropoiesis.

## Introduction

Red blood cells (RBCs) formation from progenitor cells in bone marrow (BM) requires enormous energy supply through a stepwise metabolic switch from oxidative phosphorylation to glycolysis during erythropoiesis (Van Wijk and Van Solinge, 2005; Palis, 2014). Glucose and glutamine act as the principal nutrient source for nucleotide biosynthesis to regulate the commitment of human and murine hematopoietic stem cells (HSCs) to the erythroid lineage (Oburoglu et al., 2014). It was reported that control of erythroid differentiation and maturation is also fine-tuned by fatty acids and cholesterol through the regulation of metabolic shift (Liu et al., 2017; Lu et al., 2022). Nevertheless, disruption of metabolic homeostasis, such as through restriction of energy and nutrients, may in occasion induce abnormalities of glycolytic enzymes, resulting in hemolytic anaemia (Van Wijk and Van Solinge, 2005). To meet the energy demand during nutrient deprivation, ketone bodies, namely β-hydroxybutyrate (βOHB, also known as 3-hydroxybutyrate) and acetoacetate (AcAc), alternatively serve as pivotal energy fuels and signaling mediators for metabolic and functional homeostasis in extrahepatic tissues (Robinson and Williamson, 1980; Rojas-Morales et al., 2016; Puchalska and Crawford, 2017). Nonetheless, metabolic regulations of ketone bodies in erythropoiesis under variable nutritional states remain largely understudied.

Generally, ketogenesis is initiated within hepatic mitochondrial matrix which produces ketone bodies through fatty acid oxidation catalyzed by hydroxymethylglutary-CoA synthase 2 (HMGCS2), a liver-specific ketogenic enzyme (McGarry and Foster, 1980). Subsequently, these ketone bodies are then transported from the liver into the blood circulation. Circulating concentrations of ketone bodies in healthy adults after exercise or prolonged fasting ranges between 1 mM to over 5 mM (George, 2006). In pathophysiological state, ketone body concentration can rise up to 25mM or more, such as in diabetic ketoacidosis (Laffel, 1999). On the other hand, fasting or ketogenic diets (KDs) that restrict carbohydrates and are extremely high in fat are being actively considered as an alternative to elevate circulating ketones and have potential for improving overall health and therapeutic applications (Longo and Mattson, 2014; Puchalska and Crawford, 2017). On the other hand, long-term consumption of KDs can induce side effects such as anemia especially in young women, cancer patients and in animal models (Tóth and Clemens, 2017; Arsyad et al., 2020; Hayashi et al., 2022). The mechanisms linking ketone bodies to effective erythropoiesis are yet to be well clarified.

Early erythroid progenitors are effectively dependent on erythropoietin (EPO) to support terminal erythroid differentiation, in which colony-forming unit-erythroid cells (CFU-E) subsequentially differentiate into proerythroblasts and basophilic, polychromatic, and orthochromatic erythroblasts (Haase VH. 2010). However, EPO is not the only regulator of erythropoiesis as evidenced by the failure of many cases of acute or chronic anemia to respond to EPO treatment or stimulation (Weiss and Goodnough, 2005; Thomas, 2007). Of late, several studies have reported that nutrients and histone modifications can indeed regulate erythropoiesis (Malik et al., 2017; Myers et al., 2020; Wang et al., 2021). Particularly, histone deacetylase (HDAC) was recognized to play essential roles in modulating early erythropoiesis (Vong et al., 2022). Ketone bodies mediate cell signaling not only as a metabolite participating in mitochondrial and cytoplasmic metabolic pathways (Newman and Verdin, 2014; Puchalska and Crawford, 2017), but also as an endogenous inhibitor of class I HDAC (Shimazu et al., 2013). Therefore, it is essential to clarify the roles of systemic ketone bodies in erythropoiesis, as long-term nutrient deprivation or raised ketone bodies poses challenges for healthy blood cells formation.

In this study, we have induced ketogenesis by fasting mice for 48 hours in wild-type (WT) and hepatic *Hmgcs2* KO mice, which show insufficient ketogenesis. Metabolite profiling of blood serum showed enriched fatty acid metabolism and cholesterol synthesis in fasting *Hmgcs2* KO mice compared to control mice where glycolysis is the main metabolic pathway involved in ad libitum feeding condition. We found that systemic ketogenesis has a significant effect on erythropoiesis, since in general *Hmgcs2* RNA expression is relatively low in erythrocytes and HSC. Furthermore, transcriptional study showed significantly activated fatty acids synthesis and mevalonate pathway in proerythrocytes of fasting *Hmgcs2* KO mice. Inhibition of these two pathways significantly impaired differentiation of erythroid progenitors *in vitro*. Additionally, βOHB treated erythroid progenitors showed impaired erythroid differentiation along with increased histone acetylation. Moreover, fasted *Hmgcs2* KO mice showed faster recovery of red blood cells from haemorrhage anemia after bleeding. Overall, our findings suggested that erythroid differentiation and maturation benefit from fatty acid and cholesterol synthesis in the absence of ketone bodies during nutrient deprivation and stressed erythropoiesis.

## Results

### Erythroid maturation is enhanced in ketone-less mouse under fasting condition

Our previous results have revealed that adult hepatic Alb-Cre; *Hmgcs2*^f/f^ (thereafter, KO) mouse model showed insufficient ketogenesis during fasting state, which was reflected by a significantly decreased ketone body βOHB levels in the blood serum when compared with WT mice (Arima et al., 2021). To broadly assess effects of ketogenesis on the hematopoietic system, we firstly fasted 8∼12-week-old mice for 48 hours (Longo and Mattson, 2014; Cheng et al., 2014). Both KO and WT mice showed similar blood glucose and low βOHB levels under free feeding condition (Fig. 1A). Under fasting condition, blood glucose levels were dramatically decreased and were comparable between KO and WT (Fig. 1A). However, fasting failed to induce sufficient ketogenesis in KO mice, while WT mice showed the significant increase in β OHB levels (Fig. 1A). We then refed mice for another 2 days (RF2) and 5 days (RF5) and analyzed peripheral blood (PB) from mouse tail at each time point accordingly (Fig. 1B). Strikingly, we observed a significant increase in RBCs counts and hemoglobin (Hb) levels in KO mice after 48 hours of fasting (Fig. 1C) and then recovered to the normal levels similar to WT mice after re-feeding for 2-5 days, which incidentally confirmed a previous observation (Fruhman, 1966). From this observation, we hypothesized that shifted metabolites and nutrition may play a role in regulating erythroid differentiation. Next, we analyzed bone marrow (BM) cells from fed, fasted and refed conditions respectively by flow cytometry (Fig. S1A and 1B). Erythroid differentiation and maturation can be assessed by evaluating the presentation of the cell surface markers, CD44 and Ter119 (Chen et al., 2009). Interestingly, proerythroblasts (I, ProE) were decreased under fasting condition (Fig. 1D and S1C). While we did not observe a significant difference between KO and WT for the basophilic erythroblasts (II, Baso) (Fig. 1E and S1D), polychromatic erythroblasts (III, Poly) (Fig. 1F and S1E) and orthochromatic erythroblasts (IV, Ortho) (Fig. 1G and S1F), a significant increase can be seen in maturated reticulocytes/red blood cells (V, Reti) in KO mice under fasting condition and RF2 (Fig. 1H and S1G). The frequency of all the erythroid fractions in both WT and KO mice showed recovery tendency after returning to feeding state (Fig. 1D-H). Overall, our results suggested that erythroid maturation is more enhanced in ketone-less mouse under fasting condition.

**Figure 1:**
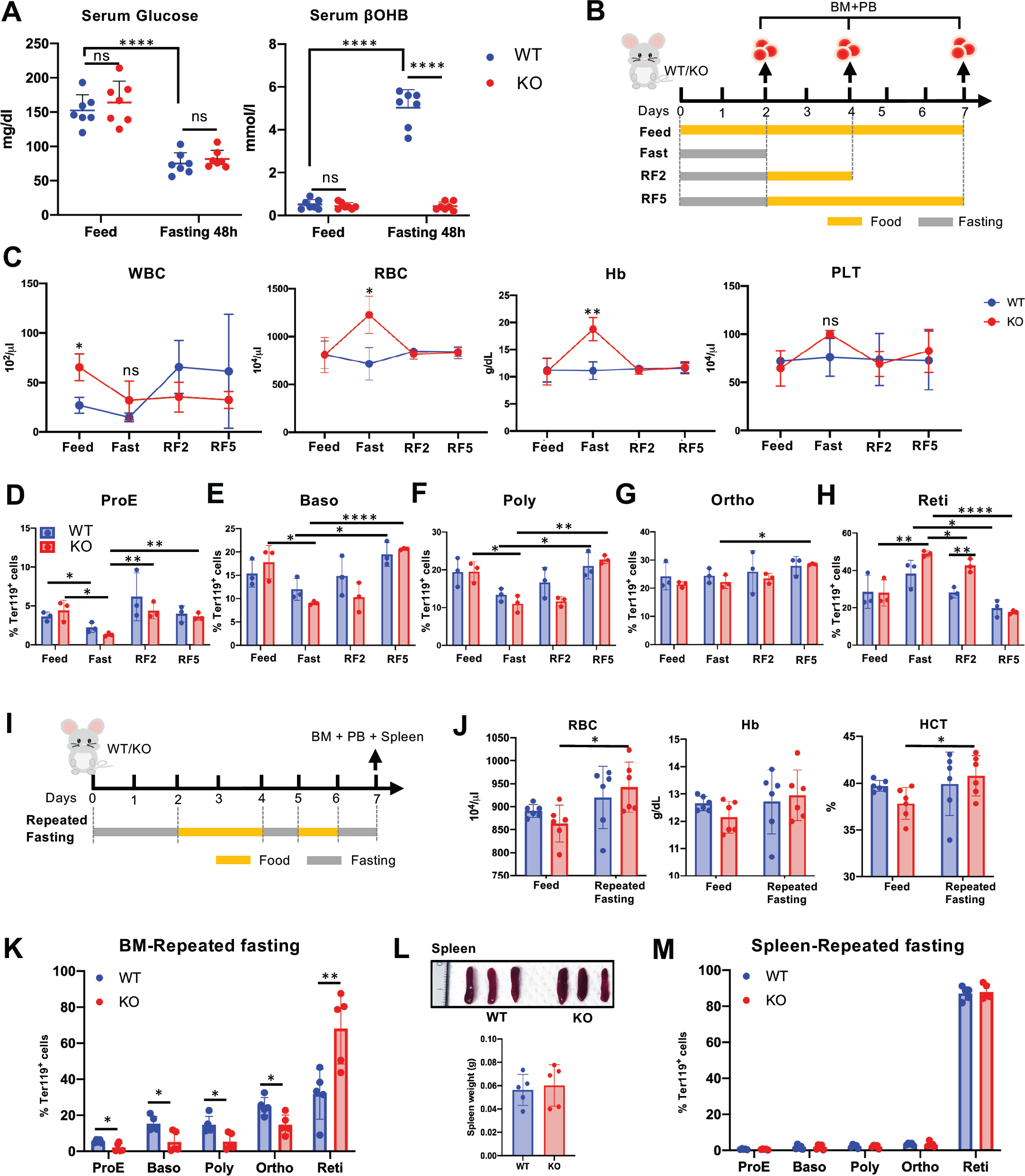
Erythroid maturation is enhanced in ketone-less mouse under fasting condition. (A) Blood glucose and ketone body (βOHB) concentration in peripheral blood (PB) of control (WT) and hepatic Alb-Cre; *Hmgcs2*^f/f^ (KO) mice after 48 hours (48h) fasting. N=7 (B) Scheme of experimental design for feeding, fasting and refeeding. Bone marrow (BM) and PB were analysed at day 2, 4 and 7 respectively. RF2 = Refeed for another 2 days after 48h fasting. RF5 = Refeed for another 5 days after 48h fasting. (C) Analysis of PB samples from WT and KO mice. WBC, white blood cells; RBC, red blood cells; Hb, haemoglobin; PLT, platelets. (D-H) Quantification of FACS analysis showing frequency of proerythroblasts (ProE) (D), basophilic erythroblasts (Baso) (E), polychromatic erythroblasts (Poly) (F), orthochromatic erythroblasts (Ortho) (G), reticulocytes (Reti) (H) in Ter119^+^ population respectively. N=3 (I) Scheme of experimental design of repeated fasting for 1 week. (J) Analysis of peripheral blood samples from WT and KO mice after 1-week repeated fasting. (K) Erythroid differentiation in BM was shown by frequency in Ter119^+^ population. N=5 (L) Spleen size and weight after 1-week repeated fasting. (M) Erythroid differentiation in spleen was shown by frequency in Ter119^+^ population. N=5. Blue bar chart indicates WT and red bar chart indicates KO. Results were shown by mean ± SD with Student’s *t* test; statistical significance was shown by ns, *P* > 0.05; *, *P* < 0.05; **, *P* < 0.01; ***, *P* < 0.001; ****, *P* < 0.0001.

### Prolonged fasting increases mature red blood cells in bone marrow but not in spleen

Taking into consideration that 48-hour fasting may introduce transient effects on mice, we modified the experimental protocol with the aims to further confirm the effect of βOHB on erythropoiesis but avoiding the harmful effects caused by ketoacidosis by adding two additional fasting cycles after 2-day refeeding and prolonging the total fasting period to 1 week (Fig. 1I, S1H). PB showed similar RBC, Hb and HCT in WT and KO mice under repeated fasting condition, while compared with feeding state KO mice showed more increased RBC and HCT (Fig.1J). Moreover, we observed significantly reduced frequency of ProE, Baso, Poly and Ortho fractions but raised maturated RBC in BM of KO mice (Fig. 1K and S1I). We hypothesized that prolonged fasting stimulates more immature erythrocytes to differentiate in ketone-less KO mice. It is known that extramedullary erythropoiesis can occur in the spleen (SPL) in response to stress (Mende and Laurenti, 2021), so we next checked the SPL to evaluate the effect of prolonged fasting on the SPL. Our results showed that the spleen size and weight were comparable between KO and WT after 1-week repeated fasting (Fig.1L). Furthermore, there was no significant difference observed from each SPL erythroid fractions between KO and WT mice (Fig. 1M). Results suggested that prolonged fasting can facilitate more mature red blood cells in the bone marrow but not in the spleen under low systemic βOHB levels.

### Less ketone body induces fatty acid synthesis under fasting condition

To investigate the βOHB-mediated metabolic regulation of erythropoiesis, we collected blood serum from feeding and 48-hour fasting mice for metabolome analysis (Sup. 2A). Under the feeding condition, glycolytic metabolites were extremely enriched in both KO and WT mice (Fig. 2A and Sup. 2B). Under the fasting condition, ketone metabolites were markedly enriched in WT mice (Fig. 2A), confirming the previous observations (Arima et al., 2021). In contrast, we observed an accumulation of serum fatty acids (FAs) in KO mice (Fig. 2A and 2B), since the liver failed to take up and convert circulating FAs into ketone bodies (Puchalska and Crawford, 2017). Additionally, in terms of FAs synthesis, this significant difference between KO and WT mice can only be detected under fasting condition (Fig. 2C and 2D, Sup. 2C), which may suggest the essential role of FAs synthesis beneath insufficient ketogenesis. Next, we aimed to investigate how systemic βOHB and FAs regulate erythroid metabolism under feeding and fasting conditions. As the mitochondrion is the key organelle for ketone body and fatty acid metabolism, we initially investigated the mitochondrial functions in erythroid populations under feeding and 48-hour fasting condition. Our observations indicated a gradually decreasing ATP contents (Fig. 2E) and mitochondrial ROS production (Fig. 2F) during erythroid differentiation and maturation. Nevertheless, we could not detect any significant difference in the metabolism of each erythroid fraction between KO and WT (Fig. 2E and 2F), suggesting that metabolic activity within cytoplasm may take place during terminal erythroid differentiation. Overall, our metabolome analysis showed that fasting condition provokes an increase in systemic βOHB levels and failure of ketogenesis might induce enrichment of fatty acid synthesis to compensate for the nutrient deprivation, which affects erythroid differentiation and maturation.

**Figure 2:**
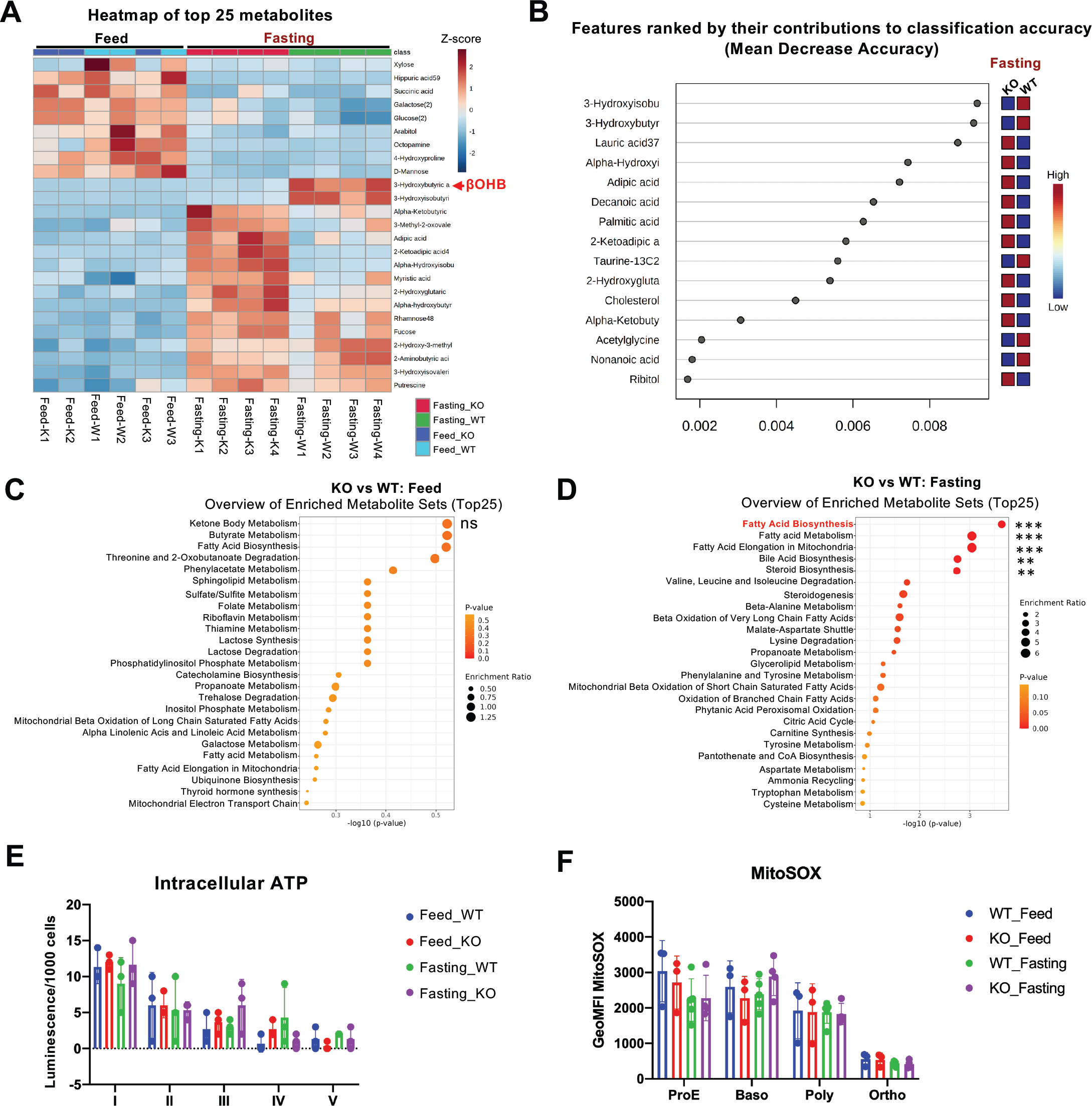
Less ketone body induces enrichment of fatty acid synthesis under fasting condition. (A) Heatmap of top 25 metabolites from mouse blood serum by metabolome analysis. Feed condition, N=3; Fasting 48h-condition, N=4. (B) Top ranking of metabolic features of blood serum in fasting condition were shown by Mean Decrease Accuracy. Overview of enriched metabolite sets were shown by dot plot under feed (C) and fasting (D) condition respectively. (E) Intracellular ATP content in erythrocytes was shown by luminescence per 1000 sorted cells for each erythroid fraction. N=3. (F) Mitochondrial ROS level for each erythroid fraction was shown by MitoSOX with geometric mean of red/green ratio. N=3. All quantified results were shown by mean ± SD with Student’s *t* test. Statistical significance was shown by ns, *P* > 0.05; *, *P* < 0.05; **, *P* < 0.01; ***, *P* < 0.001.

### Metabolic shift occurs during erythroid differentiation

To clarify the mechanisms on how ketone metabolism and fatty acid synthesis influences erythropoiesis under fasting condition, we first looked into the expression profile of the genes encoding for enzymes involved in essential metabolic pathways. In short, cell proliferation and differentiation require enormous energy supply and these energy resources are derived from complex metabolic activities such as the tricarboxylic acid (TCA) cycle within the mitochondrion as well as fatty acid synthesis and mevalonate pathway within the cytoplasm (Fig. 3A) (Granchi, 2018; Martínez-Reyes and Chandel, 2020). Using a public RNA-seq dataset of murine erythroblasts (GSE53983) (An et al., 2014), we looked for differentially expressed genes (DEGs) between the Baso, Poly and Ortho populations against the ProE population followed by a pathway analysis (Fig. S3A-C). Interestingly, we observed a distinct metabolic shift from immature erythroblasts to matured erythroblasts (Fig. 3B and Sup.3C). We selected genes of interest which encode for key enzymes that are involved in the metabolic shift from glycolysis and fatty acid metabolism to mevalonate pathway during erythroid differentiation (Fig. 3B) (Richard et al., 2019; Nemkov et al., 2022). Notably, *Hmgcs2* expression level was relatively low in ProE and further reduced during the late stages of erythropoiesis (Fig. S3D). Furthermore, glycolytic metabolism was significantly reduced in Baso, Poly and Ortho populations, indicated by decreased expression levels of *Pdk1* and *Slc25a1* (Fig. S3D). Additionally, *Oxct1*, the gene encoding succinyl -CoA:3-oxoacid CoA transferase (SCOT) which is an important enzyme that assists mitochondria to utilize external βOHB (Cotter et al., 2013), showed decreased expression levels in during erythroid differentiation (Fig. S3D). Likewise, genes related to fatty acid synthesis which are regulated by *Srebf1*, such as *Acly*, *Acaca*, *Fasn* and *Scd1*, were down regulated in terminal erythroid differentiation (Fig. S3E). Thus, we hypothesized that systemic ketone bodies and fatty acids might be an important energy source for erythroid progenitors during the early stages of erythropoiesis. Remarkably, mevalonate pathway-related genes showed significantly elevated expression levels in the late stages of erythropoiesis (Fig. S3F), suggesting the critical role of cholesterol synthesis in erythroid maturation. Overall, metabolic shift from the mitochondria to the cytoplasm is an important event for erythroid differentiation and maturation.

**Figure 3:**
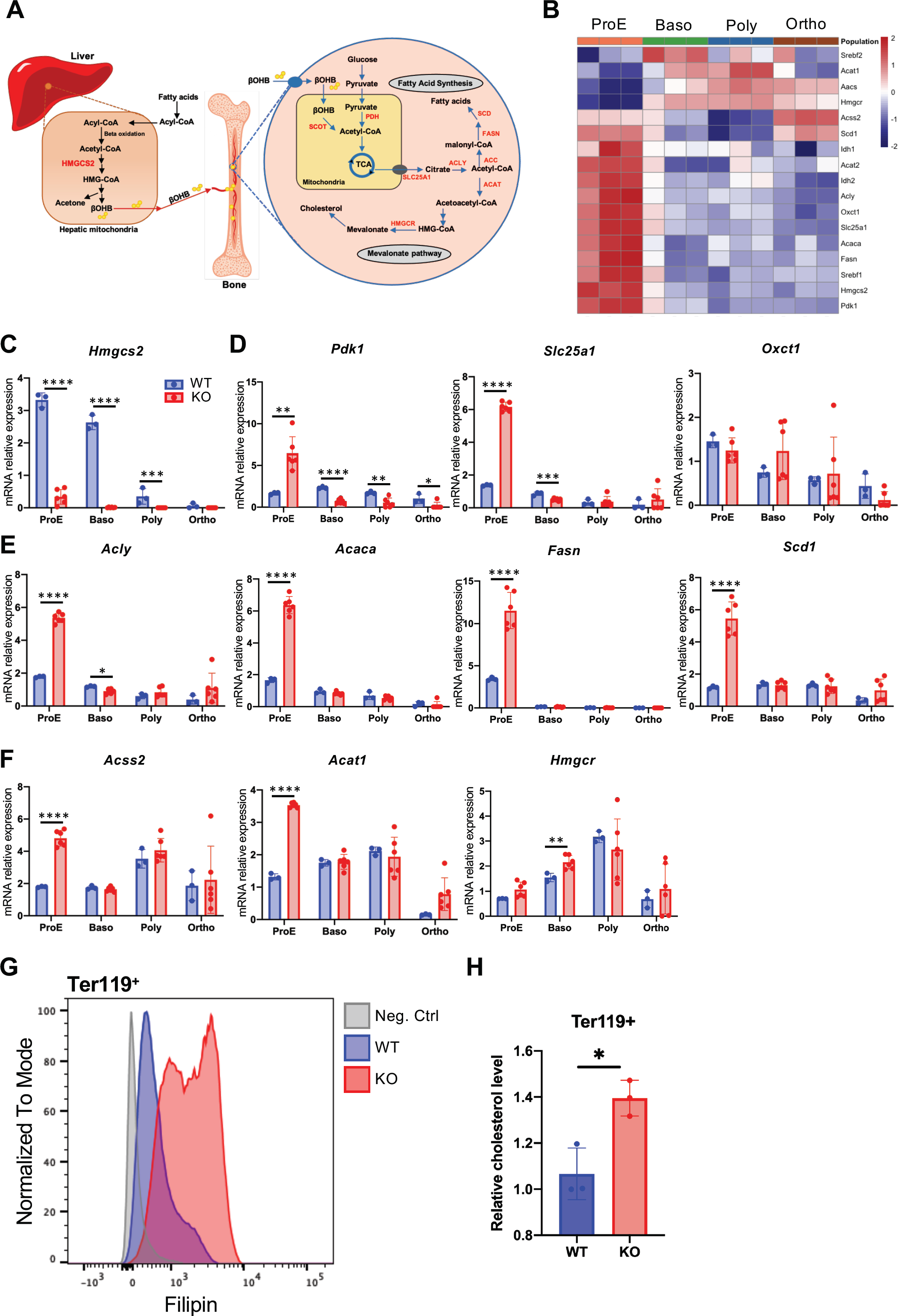
Less ketone body highly activates mevalonate pathway in erythroid differentiation and maturation during fasting state. (A) Schematic illustration of cellular metabolism in hepatic mitochondria and bone marrow cells. Under fasting condition, hepatocytes take up and convert circulating FAs into ketone bodies βOHB within hepatic mitochondria. Then, βOHB is transported into blood vessels and circulated into BM cells. Subsequently, BM cells convert βOHB and glucose into acetyl-CoA for TCA cycle and produce energy and metabolites within mitochondria, while fatty acid synthesis and mevalonate pathway take place within cytoplasm. Black arrows indicate metabolic reactions in hepatocytes; red arrows indicate βOHB circulation; blue arrows indicate metabolic reaction in BM cells. (B) RNA-seq database of normal murine erythroid populations was analysed and shown by a heatmap, indicating significant changes to genes encoding key enzymes shown in (A) during erythropoiesis. (C-F) mRNA expression levels of *Hmgcs2* (C), mitochondrial metabolism related genes (D), fatty acid synthesis related genes (E) and mevalonate pathway related genes (F) in each sorted erythroid fraction post 1-week repeated fasting were measured by RT-qPCR. All the mRNA expression levels were relative to mean expression of *Gapdh*, a control gene. N=3-6. Blue bar chart indicates WT and red bar chart indicates KO. (G) Cholesterol level in Ter119^+^ fraction was measured by Filipin III staining and analysed by flow cytometry. (H) Quantification of cholesterol level in Ter119^+^ fraction was relative to fluorescence count of negative control (Neg. Ctrl). N=3. All quantified results were shown by mean ± SD with Student’s *t* test; statistical significance was shown by ns, *P* > 0.05; *, *P* < 0.05; **, *P* < 0.01; ***, *P* < 0.001; ****, *P* < 0.0001.

### Less ketone body activates mevalonate pathway in erythroid differentiation and maturation during prolonged fasting

To validate our observations in the RNA-seq analysis, we collected erythroid progenitor fractions from KO and WT BM after one-week repeated fasting followed by RNA extraction and RT-qPCR, as 48 hours fasting did not show significant transcriptional changes in erythroid fractions between WT and KO (Fig. S4). We observed a drastic reduction of *Hmgcs2* expression in BM erythroid populations in hepatic *Hmgcs2* KO mice (Fig. 3C), suggesting that erythroid *Hmgcs2* is highly relevant in a feedback loop to synchronize systemic ketone body levels, which is consistent with the phenomena observed in other tissues such as liver, gut and kidney during fasting (Tognini et al., 2017; Venable et al., 2022). Additionally, decreased *Hmgcs2* expression levels in both KO and WT maturing erythroblasts (Fig. 3C) confirmed the same observations in the RNA-seq analysis, indicating minor effects and demands of erythroid ketogenesis on differentiation and maturation. Besides the ketogenesis from hepatocytes during fasting, kidney is known as the secondary organ that produces ketone bodies (Owen et al., 1969). However, the amount of ketone bodies produced from the kidney is insufficient to account for systemic ketogenesis significantly during fasting (Venable et al., 2022). Furthermore, *Pdk1* and *Slc25a1* expression were higher in KO ProE, which suggested that nutrients deprivation may somehow further stimulate glycolic metabolism in ProE (Fig. 3D). On the contrary, we observed lower *Pdk1* and *Slc25a1* expression in Baso from KO mice, confirming the sharp metabolic shift during erythroid maturation (Fig. 3D). *Oxct1* expression levels were comparably reduced in KO and WT during erythroid maturation (Fig. 3D). Strikingly, expression levels of fatty acid synthesis-related genes were significantly raised in ProE from KO mice compared with WT ProE (Fig. 3E). Furthermore, reduction in the expression levels of *Acly*, *Acaca*, *Fasn* and *Scd1* from Baso to Ortho fractions further confirmed the minor effects of fatty acid synthesis on erythroid maturation (Fig. 3E). Similar to the results from RNA-seq data analyses, the mevalonate pathway was significantly activated in KO ProE, indicated by increased expression levels of *Acss2*, *Acat1* and *Hmgcr* (Fig. 3F). Consistently, we also observed increased cholesterol levels in KO Ter119^+^ population (Fig. 3G and 3H). Overall, our results suggested that less ketone body favorably boosts fatty acid synthesis and cholesterol production in ProE under fasting condition, while high levels of ketone body may impair erythroid differentiation.

### Ketone body βOHB affects differentiation of early erythroid progenitors

To investigate how ketone body control functions of fatty acid synthesis and cholesterol on erythropoiesis, we cultured erythroid progenitors from BM of wild type mice and mimicked the fasting conditions in vitro. We first cultured lineage negative BM cells supplied with simplified erythroid differentiation medium (EDM) (Yang et al., 2019) with high (4.5g/l) and low (0.45g/l) concentration of glucose (Fig. 4A). We did not observe difference between high and low glucose supplement in EDM in term of erythroid differentiation after 48 hours of culture (Fig. 4B and S5A). However, more significant impairments by additional βOHB at the early stage of erythroid differentiation (Ter119^-^CD71^+^) were observed in low-glucose EDM (Fig. 4B and 4C, S5A and 5B), which conditionally mimicked the fasting condition in WT mice *in vivo* and confirmed the previous findings. Moreover, βOHB treatment significantly increased ProE (I) and Baso (II) fractions in a dose-dependent manner, while maturing factions such as Ortho (IV) and Reti (V) were reduced accordingly (Fig. 4B and 4D, S5A and 5C). The results both *in vivo* and *in vitro* indicated the impairments by βOHB on the early stage of erythroid differentiation. Equivalently, βOHB has been shown to markedly impair human erythroid differentiation in a dose-dependent manner (Liu et al., 2017). Furthermore, we found that inhibition of fatty acid synthesis and mevalonate pathway by fatostatin and lovastatin respectively efficiently suppress the erythroid differentiation (Fig. 4E and 4F). As βOHB plays an important role in the regulation of HDAC activity (Shimazu et al., 2013) and HDACs are critical to control erythroid differentiation fate (Vong et al., 2022), we further investigated the inhibition effects of βOHB for HDAC in cultured erythroid progenitors. Results showed that 10mM βOHB induced histone hyperacetylation which was harmful to erythroid differentiation and maturation (Fig. 4G). Taken together, under fasting condition, βOHB affects early stage of erythropoiesis, when fatty acid and cholesterol are important source for erythroid differentiation (Fig. 4H).

**Figure 4:**
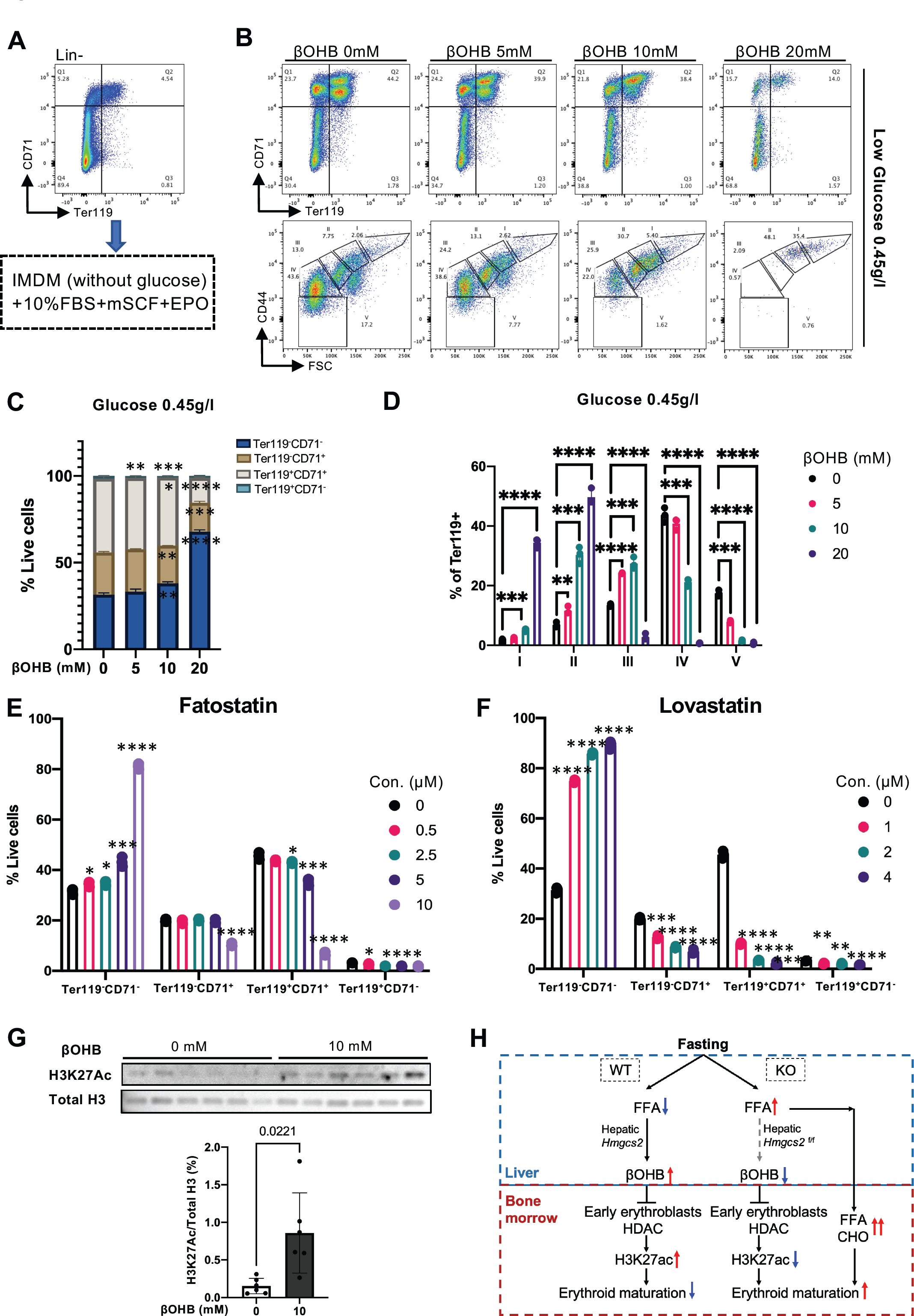
Ketone body βOHB impairs differentiation of early erythroid progenitors. (A) FACS plot showed erythroid population in lineage negative, which were collected from wide type mouse BM and cultured in erythroid differentiation medium (EDM). EDM conditions were exhibited in the black dot box. (B) FACS plots showed erythroid fractions in isolated lineage negative cells after culturing with βOHB and low glucose for 48 hours. N=3. (C) Frequency of erythroid fractions stained by CD71 and Ter119. (D) Frequency of erythroid fractions stained by CD44 and Ter119. (E) Lineage negative cells were cultured with Fatostatin in low-glucose EDM for 48 hours. Fatostatin, an inhibitor of fatty acid synthesis. N=3. (F) Lineage negative cells were cultured with Lovastatin in low-glucose EDM for 48 hours. Lovastatin, an inhibitor of mevalonate pathway. N=3. (G) Western Blot showing histone acetylation levels of early erythroid progenitors after βOHB (10mM) treatment for 48 hours. Representative experiment of N=6. (H) Scheme showing systemic ketone body effects on erythropoiesis. Blue arrows indicate downregulation and red arrows indicate upregulation. All quantified results were shown by mean ± SD with Student’s *t* test; statistical significance was shown by ns, *P* > 0.05; *, *P* < 0.05; **, *P* < 0.01; ***, *P* < 0.001; ****, *P* < 0.0001.

### Fasted mice with less ketone body recover faster from hemorrhage

Our observations above indicated that high levels of ketone body βOHB impairs erythropoiesis through complex metabolic reactions. We then next tried to ascertain the physiological role of ketone body in stressed hematopoiesis under feeding and fasting conditions. To achieve this, we induced acute anemia in KO and WT mice by serial bleeding for 3 days (Day1-3) followed by feeding or 48-hour fasting (Day3-5) (Fig. 5A). We found that the body mass of fasted mice was remarkably reduced (Fig. 5B), and the body mass of KO mice was comparable with the WT mice in both feeding and fasting conditions during and after bleeding. Moreover, bleeding caused acute anemia at day 4, indicated by a sharply decreased RBCs counts, HGB and HCT (Fig. 5C). It is known that spleen is one of the organs responding for stressed erythropoiesis (Hara and Ogawa, 1976; Chiu et al., 2015). Surprisingly, we observed significantly less splenomegaly with lower red pulp macrophages (RPMs) in KO mice after fasting (Fig. 5D-F). We then analyzed the different erythroid fractions of the BM and spleen in KO and WT mice (Fig. 5G and 5H). Under the feeding condition, BM and spleen erythropoiesis was comparable in KO and WT (Fig. 5I and J). However, under the fasting condition, KO mice showed significantly enhanced erythroid maturation in BM (Fig. 5K). Likewise, a similar observation was seen in the spleen which showed increased erythropoiesis in KO mice under fasting (Fig. 5L). Thus, it is likely that less ketone body under fasting can be beneficial for both BM and spleen erythropoiesis under the stressed conditions.

**Figure 5:**
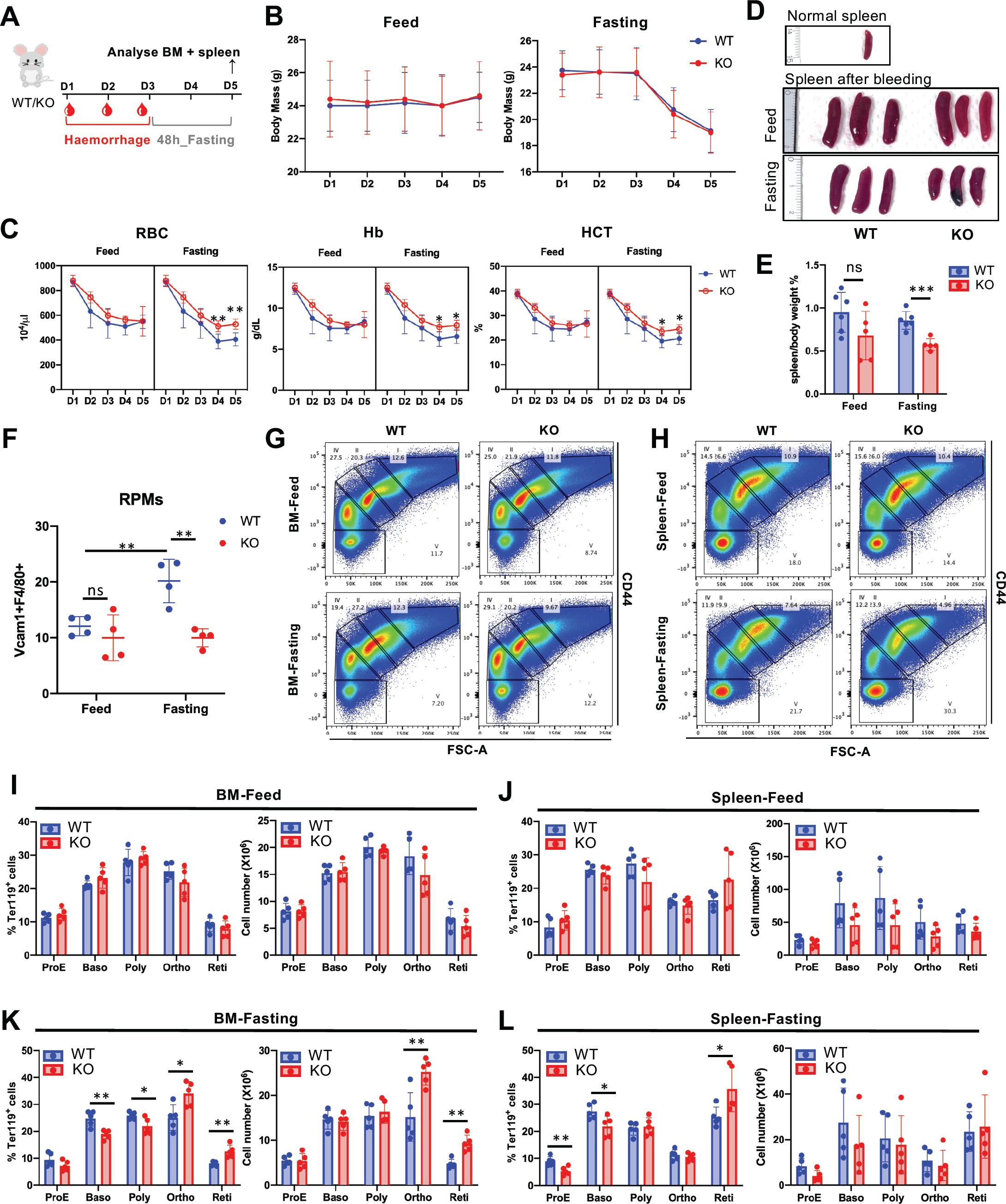
Less ketone body rescues anaemia caused by haemorrhage stress. (A) Scheme of experimental design with combination treatment of haemorrhage (bleeding) and fasting. (B) Mouse body mass was measured during bleeding and recovery phase under feed (left panel) or fasting (right panel) condition. (C) Analysis of peripheral blood samples from WT and KO mice. Anaemia was indicated by decreased RBC, Hb and HCT. N=5-6. Spleen size (D) and spleen/body weight (E) was measured after 48 hours post third bleeding (day 5). (F) Frequency of red pulp macrophages (RPMs) in spleen. N=4. (G-H) FACS plot showed erythroid differentiation and maturation in bone marrow (G) and spleen (H) under feed and fasting conditions after bleeding. (I and K) Erythroid differentiation and maturation in bone marrow were shown by frequency and cell numbers of each erythroid faction analysed by flow cytometry. (J and L) Erythroid differentiation and maturation in spleen under feed and fasting conditions after haemorrhage stress were shown by frequency and cell numbers of each erythroid faction analysed by flow cytometry. N= 5. All quantified results were shown by mean ± SD with Student’s *t* test; statistical significance was shown by ns, *P* > 0.05; *, *P* < 0.05; **, *P* < 0.01; ***, *P* < 0.001.

## Discussion

Ketone bodies can act as a guard to protect mice from chemotoxicity (Lee et al., 2010), preserve mitochondrial energy of neonatal hepatocytes (Arima et al., 2021), mediate intestinal stemness (Cheng et al., 2019) and improve immunity by inhibiting inflammation (Youm et al., 2015; Goldberg et al., 2017, 2020). However, as raised ketone bodies may pose challenges for healthy blood cells formation, it is crucial to understand the mechanisms linking ketone bodies with terminal erythroid differentiation.

Here, we established a low systemic ketogenesis animal model by conditionally knocking out *Hmgcs2* in murine hepatocytes (Arima et al., 2021). In agreement with observations in *Drosphila* (Shim et al., 2012), fasting promotes the production of matured RBCs as seen in ketone-less mice after 48 hour fasting (Fig. 1C-H), which can be reversed to normal state by refeeding (Fig. 1C-H). The reduction of erythroid differentiation and maturation in WT BM with high βOHB is potentially due to a differentiation arrest as these events are reversible after refeeding (Fig.1). Cellular differentiation requires an enormous energy supply which can be reached by increasing mitochondrial metabolism. Under fasting condition, we found that mitochondrial-related genes, such as *Pdk1* and *Slc25a1*, exhibited lower expression levels in WT ProE compared to KO (Fig. 2D), which may indicate a sign of differentiation arrest with high βOHB. Furthermore, we found that ketone-less mice under prolonged fasting for 1 week showed a further enhancement of RBCs maturation (Fig. 1I-K) but reduced frequency of hematopoietic stem and multipotent progenitor cells (HSPC) in BM, while HSPC in control mice were comparable between feeding and fasting condition (Fig. S6). This suggests that systemic ketogenesis may play a role to maintain HSC homeostasis under fasting stress, which reflects the findings from a previous study which reported that prolonged fasting can protect HSPC from stress and restore lineage balance (Cheng et al., 2014), but it is also important that how other factors such as the niche (e.g. bone marrow adipocytes) can effect HSCs during fasting condition to be taken into account (Kaastrup et al., 2021).

Maintenance of HSC homeostasis and in deciding differentiation fate during nutrient deprivation, metabolic regulation definitely plays a center role to compensate for the lack of energy source and help cellular adaptation (Nakamura-Ishizu et al., 2020; Mistry et al., 2022). Strikingly, we found that metabolic pathways are sharply shifted from mitochondria to the cytoplasm (Fig. 2 and 3), in which the mevalonate pathway is favorably activated during terminal erythroid differentiation and maturation. This metabolic shift from glycolysis to fatty acid metabolism and cholesterol synthesis is reasonable as organelles, particularly the mitochondrion, are diminished during terminal erythroid differentiation. A study by Lu et al (Lu et al., 2022) had also confirmed that cholesterol is essential for erythroid differentiation. Furthermore, we illustrated that increased fatty acid synthesis and cholesterol levels in erythrocytes of ketone-less mouse can facilitate erythroid maturation, while increased βOHB or inhibition of fatty acid synthesis and mevalonate pathway exhibited impairment of erythroid differentiation (Fig. 4A-F), which is highly consistent with the previous study on human erythroid differentiation (Liu et al., 2017).

On one hand, in terms of impairment of erythroid differentiation, we showed that βOHB administration inhibited the HDAC activity, thereby increasing the histone acetylation activity (Fig. 4H). Similarly, one study reported that βOHB reduced class I HDAC activity resulting in increased H3K27ac activity and GATA2 expression but suppressed the expression of GATA1 and hemoglobin genes (Liu et al., 2017). This contrasting expression of GATA genes is considered to be harmful for erythroid differentiation (Moriguchi and Yamamoto, 2014) as GATA1 deacetylation, which is mediated by class I HDAC, is particularly essential for terminal erythroid differentiation (Yan et al., 2021). Thus, our results suggest that fasting-induced high systemic βOHB levels suppressed erythroid differentiation partially through histone hyperacetylation in early erythroid progenitors.

However, on the other hand, under starvation stress, βOHB is a sort of response metabolite to enhance stress resistance and protect cells against highly oxidative stress and damage (Rojas-morales et al., 2020), which positively extends cellular lifespan (Sohal et al., 2009; Edwards et al., 2014). Thus, it is possible that βOHB also play a role in maintaining hematopoietic homeostasis, since our results showed that repeated fasting significantly reduced HSPCs frequency but increased matured RBC in ketone-less *Hmgcs2* KO mice. Therefore, the reduction of erythroid maturation in WT mice might be an essential step as an anti-oxidative response to maintain metabolic homeostasis in BM.

Upon exposure to hemorrhagic and fasting stress, we observed that ketone-less KO mice appear to recover more rapidly from acute anaemia caused by bleeding (Fig. 5). Moreover, fasted ketone-less KO mice have a smaller spleen size and weight in comparison to control mice. This indicated that a more efficient generation and maturation of RBCs occurs in the BM and spleen of fasted ketone-less KO mice, whereas in fasted control mice, the presence of ketone bodies impair or delay erythroid differentiation and maturation (Fig. 5). Although it is unclear whether reduction or removal of ketone bodies can be beneficial, such as for the treatment of anemia, our study demonstrated that high levels of ketone bodies in the blood can have negative effects on erythroid differentiation which we hope to make aware especially to those on a long-term ketogenic diet or starvation. More importantly, our study shed a light on multi-dimensional views from metabolism to transcriptome level to better understand erythroid differentiation dietarily, in which insufficient systemic βOHB partially reduces histone acetylation and boosts fatty acid and cholesterol to support mature RBCs formation, which could be a potential reference for dietary and therapeutic design.

## Methods

### Mouse study

Hepatocytes specific loss of ketogenesis (Alb-Cre; *Hmgcs2*^flox/flox^) mice [C57BL/6J] were generated and provided by Y.A. (Arima et al., 2021). Mice at 8 to 12 weeks of age were either fasted for 48 hours or fed ad libitum for experiments. All mice were maintained under animal care guidelines of Kumamoto University. All procedures were performed in accordance with the Kumamoto University animal care guidelines (approval reference no. A2033-073).

### Measurement of blood glucose and beta-hydroxybutyrate

Whole-blood samples were obtained from the superficial temporal vein of mice after anesthesia. Immediately, we measured the blood glucose by using ACCU-CHEK Aviva nano (Roche). We measured blood beta-hydroxybutyrate concentration by using FreeStyle Precision Neo and FreeStyle Precision Blood β-ketone test strips (Abbott).

### Complete Blood Counts

Mouse peripheral blood was obtained from the superficial temporal vein of mice after anesthesia. Blood samples were collected in K2EDTA tubes (BD), and analysed via the Celltac Alpha veterinary hematology analyzer, pocH-100i V Diff (Sysmex).

### Gas chromatography-mass spectrometry analysis

Mice were either fasted for 48 hours or fed ad libitum followed by whole-blood collection. Then mouse serum samples were obtained by centrifugation at 3,000g at room temperature (RT) for 10 min. Subsequently, samples were analyzed on a Shimadzu 50GC-MS/MS. The chromatograms and mass spectra were analyzed utilizing the GC-MS solution software v4.5 (Shimadzu). Compounds were determined with the Smart Metabolite Database v2 (Shimadzu). Metabolome results were analyzed by R and MetaboAnalyst 5.0 platform (https://www.metaboanalyst.ca/).

### Flow cytometry analysis and antibodies

Total bone marrow (BM) cells were obtained by flushing femur and tibia bones with Dulbecco’s Modified Eagle Medium (DMEM, Sigma) containing 10% fetal bovine serum (FBS, Biowest). Total cell numbers were counted after washing with cold FACS buffer (phosphate-buffered saline (PBS) with 2% FBS). BM cells were stained with fluorescence-conjugated antibodies against different cell surface markers and analyzed by flow cytometry. Following antibodies were used: c-Kit (2B8, PE-Cy7; BD Biosciences), Sca-1 (E13-161.7, FITC; eBioscience), CD150 (TC15-12F12.2, BV-421; BioLegend), CD48 (HM48-1, APC-Cy7; BioLegend), Ter119 (TER-119, PerCP-Cy5.5; BD Biosciences), CD4 (GK1.5, PerCP-Cy5.5; BD Biosciences), CD8a (53-6.7, PerCP-Cy5.5; BD Biosciences), B220 (RA3-6B2, PerCP-Cy5.5; BD Biosciences), Gr-1 (RB6-8C5, PerCP-Cy5.5; BD Biosciences), Mac-1 (M1/70, PerCP-Cy5.5; BD Biosciences). Cell population was identified as following definition: LSK (Lineage-Sca1+cKit+, LSK), MPP2 (LSK, CD48+ CD150-), MPP3 (LSK, CD48+ CD150-), ST-HSC (LSK, CD48-CD150-) and LT-HSC (LSK, CD48-CD150+). All staining samples were incubated in FACS buffer for 30min on ice following by cell sorting using Aria (BD Life Science-Bioscience) or following by flow cytometry analysis using FACSymphony (BD Life Science-Bioscience). Data were analyzed by using FlowJo software v10.

### Mitochondrial metabolism analysis

Mitochondrial superoxide production and intracellular ATP production of erythroid fractions was determined with MitoSOX Red Mitochondrial Superoxide Indicator (Thermo Fisher Scientific) and “Cell” ATP Assay reagent Ver.2 (FUJIFILM) respectively, following manufacturer’s instructions. Briefly, for intracellular ATP contents, BM cells were collected and stained with erythroid surface makers, CD44 and Ter119, as described above. Then, 1000 cells per each erythroid fraction were sorted into 96-well white plate with glass bottom. After incubating cells with the same amount of working solution for 10 min at RT, the plate was analyzed with luminescence plate reader (Synergy H1). BM cells were collected and stained with MitoSOX Red probe for 30min at 37°C following by erythroid surface markers staining for 30min on ice. All the samples were analyzed by flow cytometry to determine the fluorescence intensity of MitoSOX Red probe.

### RNA extraction and qRT-PCR

Total erythroid RNA from BM was isolated using ISOGEN (Wako) following manufacturer’s instructions. Reverse transcription was performed using a PrimeScript RT reagent Kit (TaKaRa) following manufacturer’s instruction. Gene expression was determined using Luna Universal qPCR Master Mix (BioLabs) using Roche LightCycler96 system. Data was normalized relative to *Gapdh*. Mouse primers sequence information was shown in supplementary Table 1.

### Culture of murine erythroid progenitors and inhibitors

BM cells were obtained by flushing femur and tibia bones from 8w age of wide type mice with Dulbecco’s Modified Eagle Medium (DMEM, Sigma) containing 10% fetal bovine serum (FBS, Biowest). Then, lineage-negative bone marrow cells were isolated by autoMACS as previously described. Briefly, after washing with PBS, BM cells were stained with Biotinyl lineage markers for 30 min on ice, following by incubation with magnetic microbeads conjugated streptavidin for another 20 min at 4°C. Following biotinylate antibodies were used: CD3ε (145-2C11: Bio Legend), B220/CD45R (RA3-6B2: Bio Legend), Mac-1 (M1/70: Bio Legend), Gr-1 (RB6-8C5: Bio Legend), CD4 (SK3: Bio Legend), CD8 (53-6.7: Bio Legend), Ter119 (TER-119: Bio Legend). Next, lineage negative cells were selected by Magnetic Cell Sorter AutoMACS (Miltenyibiotec), following manufacturer’s instructions. Isolated progenitor cells were cultured for 48 hours in erythroid differentiation medium (Iscove Modified Dulbecco Medium, Stemcell Technologies) supplied with 10% FBS, 10ng/ml mouse Stem Cell Factors (mSCF) (R&D-455), 2U/ml epoetin and 0.45 g/l or 4.5g/l glucose (Yang et al., 2019; Huang et al., 2018). Erythroid progenitors were cultured with βOHB (3-Hydroxybutyric acid, Sigma), fatostatin (HY-14452, MedChemExpress), an inhibitor of fatty acid synthesis or lovastatin (S2061, Selleckchem), an inhibitor of mevalonate pathway, in a dose-dependent manner. After 48 hours, cells were stained with erythroid surface markers and analyzed by flow cytometry.

### Measurement of cholesterol level

To measure free cholesterol level in erythrocytes, BM cells were stained with erythroid surface markers and Filipin III (Sigma, SAE0087) as previously described (Muller et al., 1984). Briefly, BM cells were fixed with 4% paraformaldehyde for 1 hour at RT. Then, glycine was added to quench the paraformaldehyde following by Filipin (50μg/ml) staining for 30 min at RT. After washing cells with PBS, cells were stained with CD44 and Ter119 for 30 min on ice following by analysis with flow cytometry. Filipin fluorescence intensity was analyzed with FlowJo software ver.10.

### Histone protein extraction and Western Blot

We used a FACS Aria III for cell sorting as described above and collected immature erythroid cells (2×10^5^) after *in vitro* culture with βOHB for 48 hours. Collected cells were lysed with extraction buffer (0.1M Tris-HCl, 150mM NaCl, 1.5mM MgCl_2_, and 0.65% NP40) and centrifuged at 12,000 rpm for 1 minute. Pellets were incubated with 0.2M H_2_SO_4_ and acid extracts were used for western blotting analysis.

Western Blot was performed with an SDS-PAGE system. Blotted gels (Bio-Rad, TGX Stain-Free^TM^ FastCast^TM^ Acrylamide Solutions) were transferred to nitrocellulose membranes using the Trans-Blot Turbo Transfer System (Bio-Rad). Membranes were immersed in 5% skim-milk containing blocking solution and reacted with specific primary antibodies. Mouse derived anti-histone3 antibody (Abcam, ab24324) and rabbit derived anti-H3K27Ac antibody (Cell Signaling, 8173S) were used. For fluorescent western blotting, goat anti-rabbit IgG IRDye 800CW (LI-COR Bioscience Systems) and anti-mouse IgG IRDye 600RD (LI-COR Bioscience Systems) were used as the secondary antibody. Images were obtained by Image analyzer (Bio-Rad, ChemiDoc Touch).

### Serial blood withdrawal and fasting

To generate acute anemic stress, 300-400 μl of peripheral blood was obtained from the submental vein of the mice (8-12 weeks old) for three consecutive days following another 48 hours fasting. They were euthanized via isoflurane inhalation 5 days after first bleeding and subsequently subjected to FACS analysis.

### RNA-sequencing data analysis

Count matrices generated from RNA-seq data in four stages of murine erythropoiesis (An et al., 2014) were obtained from the Gene Expression Omnibus (GEO) accession GSE53983 and analyzed using edgeR. Data was normalized and converted to counts per million (CPM) followed by batch effect removal using the replicate groups as the source of batch effect. Significance was determined using one-way ANOVA followed by post hoc Tukey’s test and defined as a P value of less than 0.05.

### Statistics

Experiments were performed more than three times. Results are expressed as the mean ± SD. Three-group comparisons were analyzed by one-way ANOVA and Holm-Sidak’s multiple-comparisons test. Two-group comparisons were carried out with the Welch’s two-sided t-test. Statistical analyses were performed with Prism 8 (GraphPad).

### Online supplemental material

Supplementary Figure 1 shows erythroid maturation is enhanced in ketone-less mouse under fasting condition. Supplementary Figure 2 shows metabolome analysis of blood serum after 48 hours fasting. Supplementary Figure 3 shows metabolic shift occurs during erythroid differentiation. Supplementary Figure 4 shows metabolic genes involved in erythroid differentiation and maturation after 48 hours fasting. Supplementary Figure 5 shows ketone body βOHB impairs early erythroid progenitor differentiation. Supplementary Figure 6 shows frequency of HSPCs was reduced in ketone-less mouse after 1-week repeated fasting. Supplementary Table 1 shows mouse primers sequence information for quantitative PCR.

## Acknowledgements

We thank Miho Kataoka, Takako Ideue and Alban Johansson for technical assistance; Dr. Tatsuya Morishima for kindly providing Epo as a gift and active discussion on *in vitro* erythroid cultures; Dr. Kiyoka Saito for kindly providing fatty acids as gifts and active discussion on *in vitro* erythroid cultures; Dr. Benjy Tan Jek Yang for assistance of bioinformatic analysis. Toshio Suda is supported by the Singapore Translational Research Investigator Award from the National Medical Research Council in Singapore (NMRC/STaR 18 may-0004) and in part by a JSPS KAKENHI Grant-in-Aid for Scientific Research (S; grant no. 26221309).

## Author contributions

WM performed all the experiments and data analysis. YA and YX kindly generated and provided hepatic *Hmgcs2* KO mouse. TY provided technical training and guidance on flow cytometry and cell sorting. WM analyzed erythroid bulk RNA-seq data with technical assistance from JYT. Moreover, WM performed metabolome analysis with the assistance and advise from TU and YA. KM provided guidance and advice for erythroid culture *in vitro*. TS, YA, YT and TU provided guidance and advice for experimental design and results interpretation. WM and TS conceived the project, designed experiments, interpreted results and wrote the manuscript. All authors read and approved the final manuscript.

