## Supplemental Files for "Metabolic regulation in erythroid differentiation by systemic ketogenesis in fasted mice"

**Supplementary Figure 1: Erythroid maturation is enhanced in ketone-less mouse under fasting condition.** (A) Gating strategy of FACS analysis to show erythroid differentiation and maturation. Purple arrow indicates erythroid differentiation direction. (B) FACS plots showed frequency of each erythroid fractions in $Ter119^{+}$ population in WT and KO at different time point respectively. (C-G) Cell number of each erythroid fraction. N=3. (H) Blood glucose and βOHB concentration were shown during 1-week repeated fasting. (I) Representative FACS plots of erythroid differentiation and maturation after 1-week repeated fasting. All quantified results were shown by mean ± SD with Student's *t* test; statistical significance was shown by ns, $P > 0.05$; *, $P < 0.05$; **, $P < 0.01$.

**Supplementary Figure 2: Metabolome analysis of blood serum after 48 hours fasting.** (A) PCA plots of blood serum metabolome analysis by MS-Spectra. (B) Overview of enriched metabolite sets were shown by dot plot under feed and fasting condition respectively. (C) Relative abundance of fatty acid synthesis related metabolites under fasting condition.

**Supplementary Figure 3: Metabolic shift occurs during erythroid differentiation.** (A) PCA plot of RNA-seq datasets. (B) Heatmaps of RNA-seq datasets showed up and down regulated differentially expressed genes in 4 erythroid fractions. (C) Pathway analysis of up and down regulated genes shown in (B). (D-F) RNA expression levels of *Hmgcs2*, mitochondrial metabolism related genes (D), fatty acid synthesis related genes (E) and mevalonate pathway related genes (F) in each erythroid fraction were shown by box graph respectively. N=3. Red box indicates ProE, green indicates Baso, blue indicates Poly and purple indicates Ortho. All quantified results were shown by mean ± SD with 1 way ANOVA and post-hoc Tukey's test; statistical significance was shown by ns, $P > 0.05$; *, $P < 0.05$; **, $P < 0.01$; ***, $P < 0.001$; ****, $P < 0.0001$.

**Supplementary Figure 4: Metabolic genes involved in erythroid differentiation and maturation during 48 hours fasting**. (A-B) Mitochondrial metabolism related genes, (C-F) fatty acid synthesis related genes and (G-H) mevalonate pathway related genes in each sorted erythroid fraction post 48 hours fasting were measured by RT-qPCR. All the mRNA expression levels were relative to mean expression of *Gapdh*, a control gene. N=5-6. All quantified results were shown by mean ± SD with Student's *t* test; statistical significance was shown by ns, $P > 0.05$.

**Supplementary Figure 5: Ketone body βOHB impairs early erythroid progenitor differentiation.** (A) FACS plots showed erythroid fractions in isolated lineage negative cells after culturing with βOHB and high glucose (4.5g/l) for 48 hours. (B) Frequency of erythroid fractions stained by CD71 and Ter119. (C) Frequency of erythroid fractions stained by CD44 and Ter119. N=3. All quantified results were shown by mean ± SD with Student's *t* test; statistical significance was shown by ns, $P > 0.05$; *, $P < 0.05$; **, $P < 0.01$; ***, $P < 0.001$; ****, $P < 0.0001$.

**Supplementary Figure 6: Frequency of HSPCs was reduced in ketone-less mouse after 1-week repeated fasting**. (A) Representative FACS plots of HSPCs. (B-E) Frequency of HSPCs were decreased in KO mice after 1-week repeated fasting. N=3. All quantified results were shown by mean ± SD with Student's *t* test; statistical significance was shown by ns, $P > 0.05$; *, $P < 0.05$.

### Supplementary Figure 1

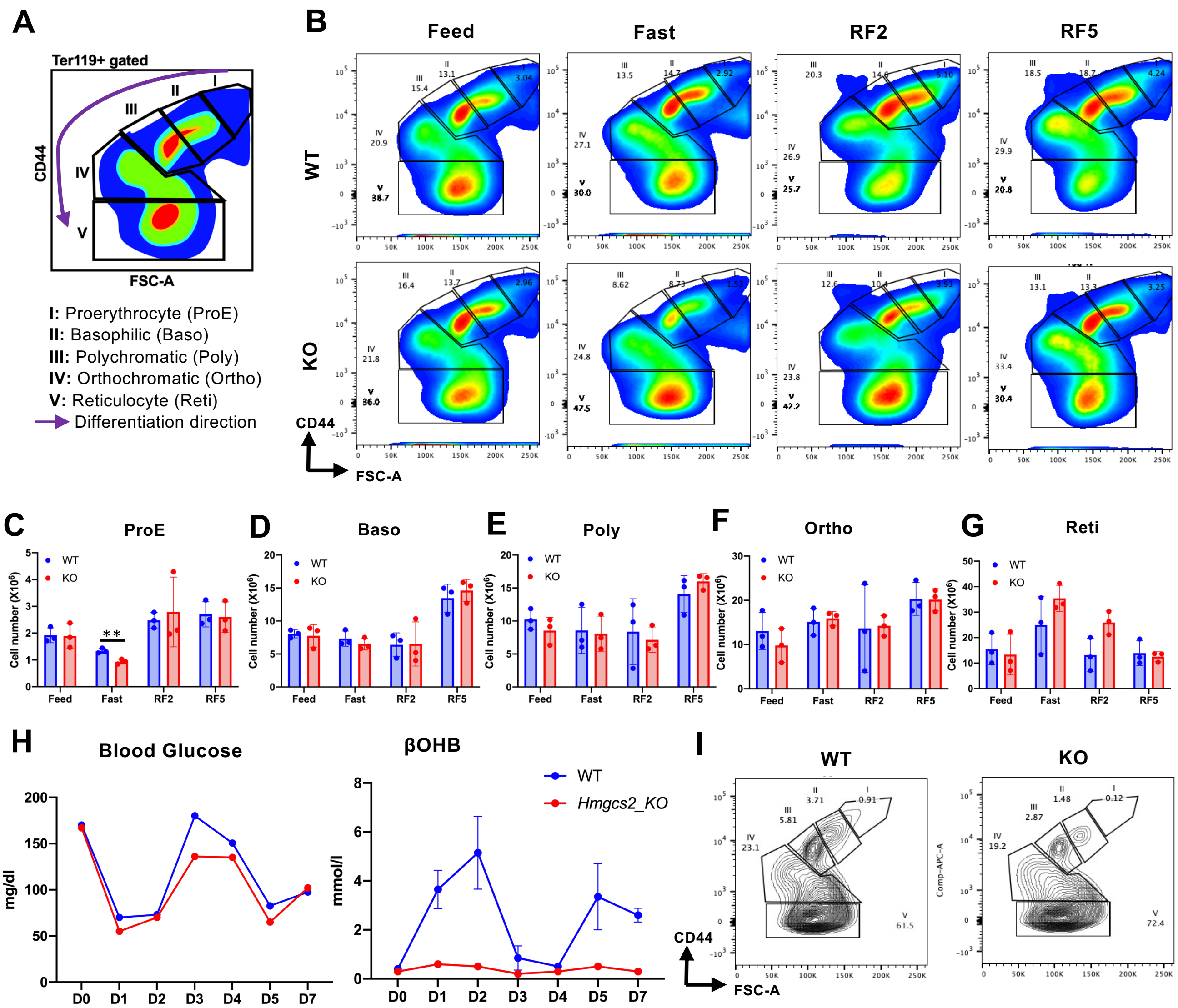

### Supplementary Figure 2

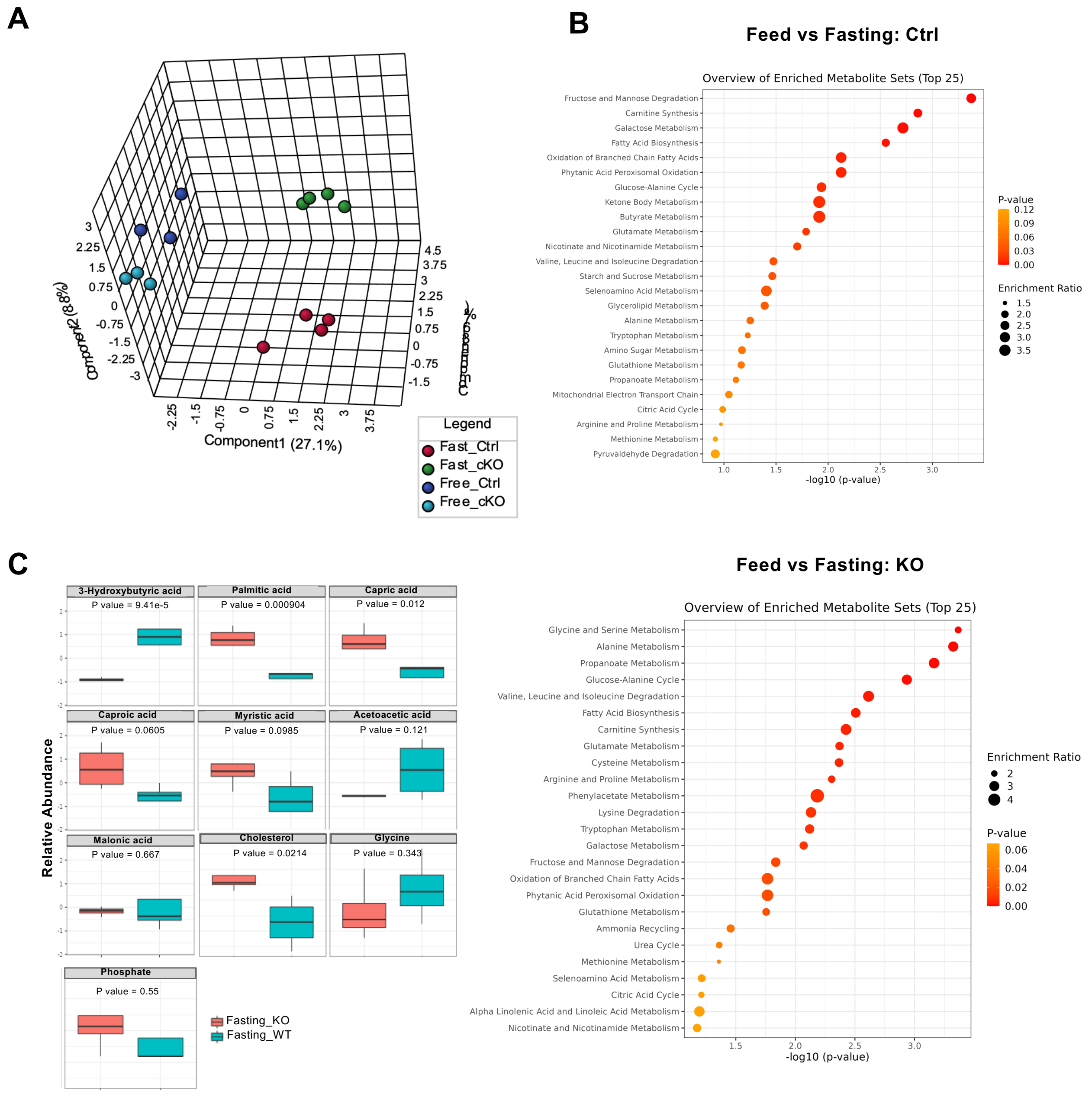

### Supplementary Figure 3

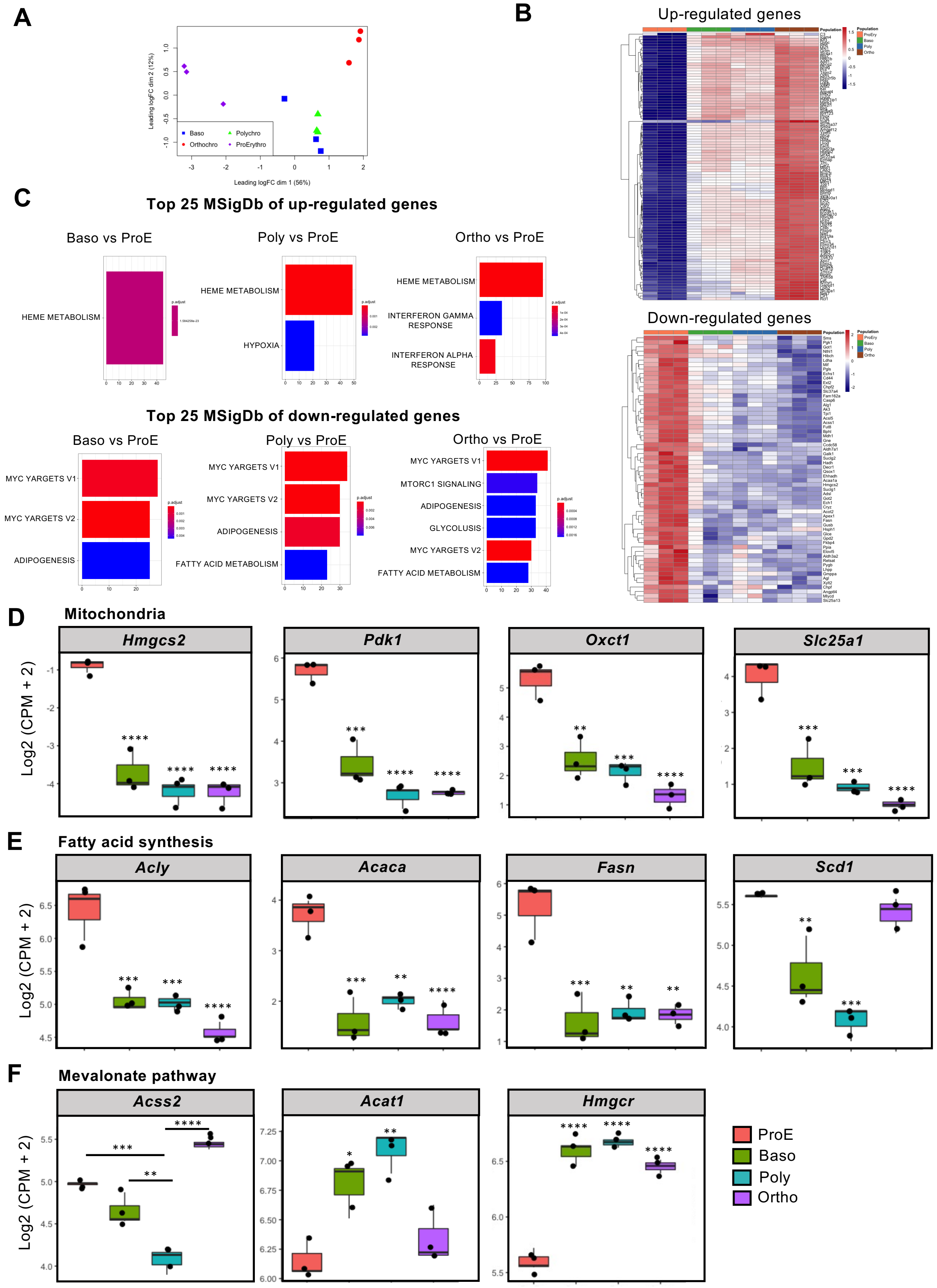

### Supplementary Figure 4

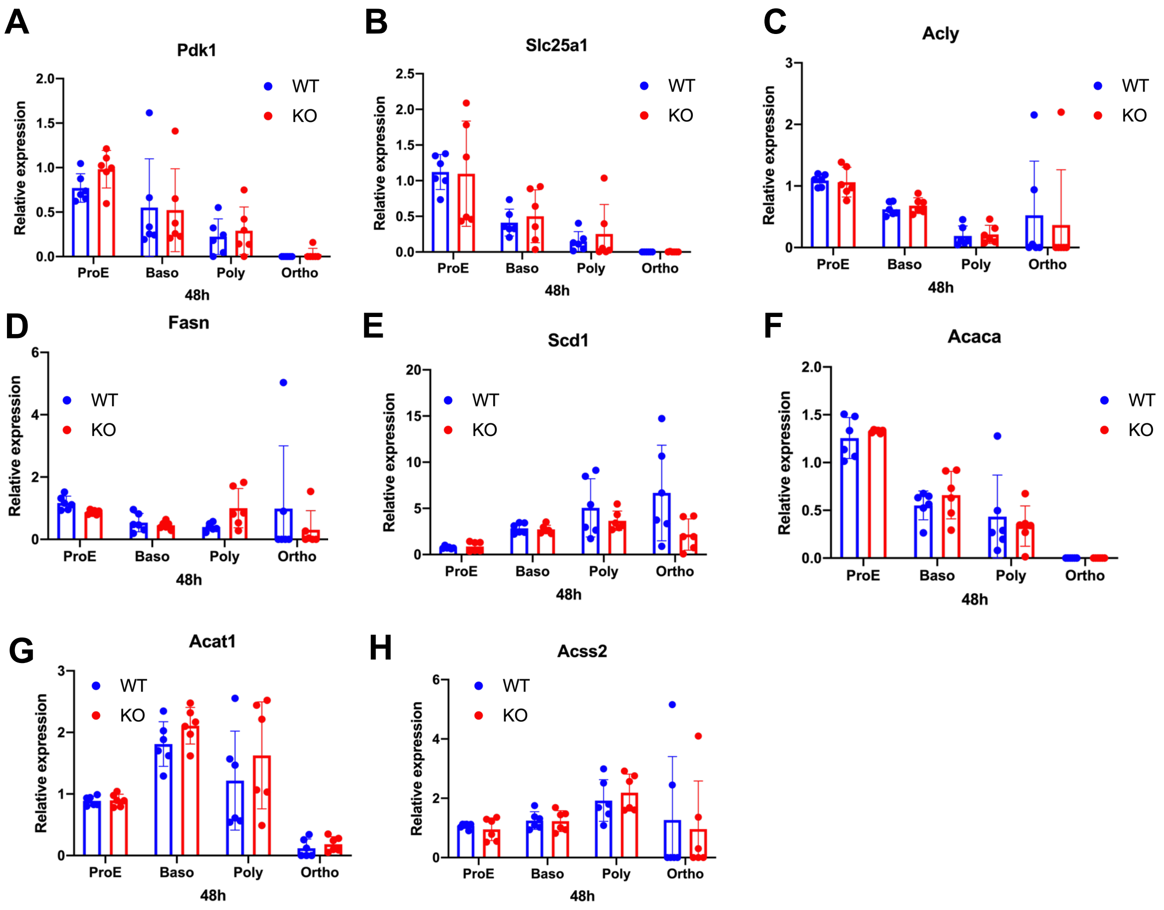

### Supplementary Figure 5

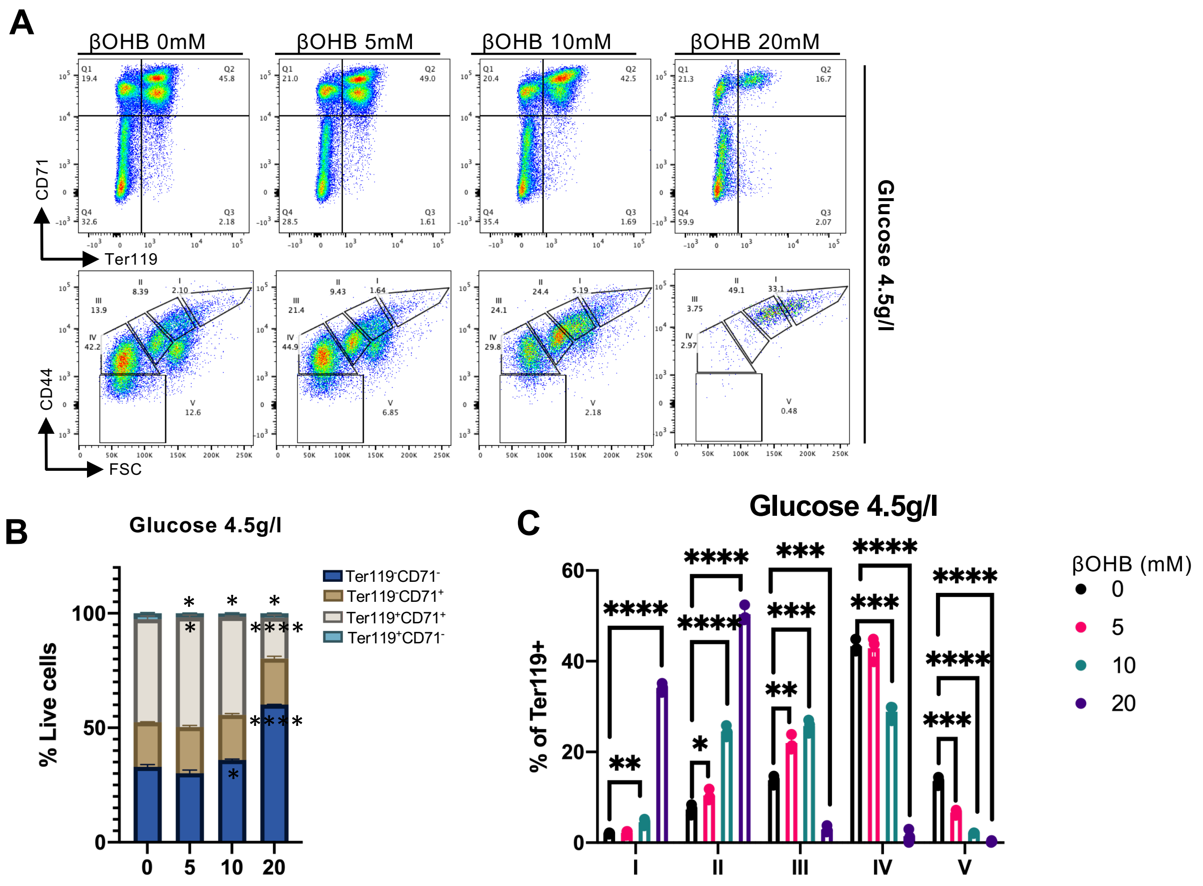

### Supplementary Figure 6

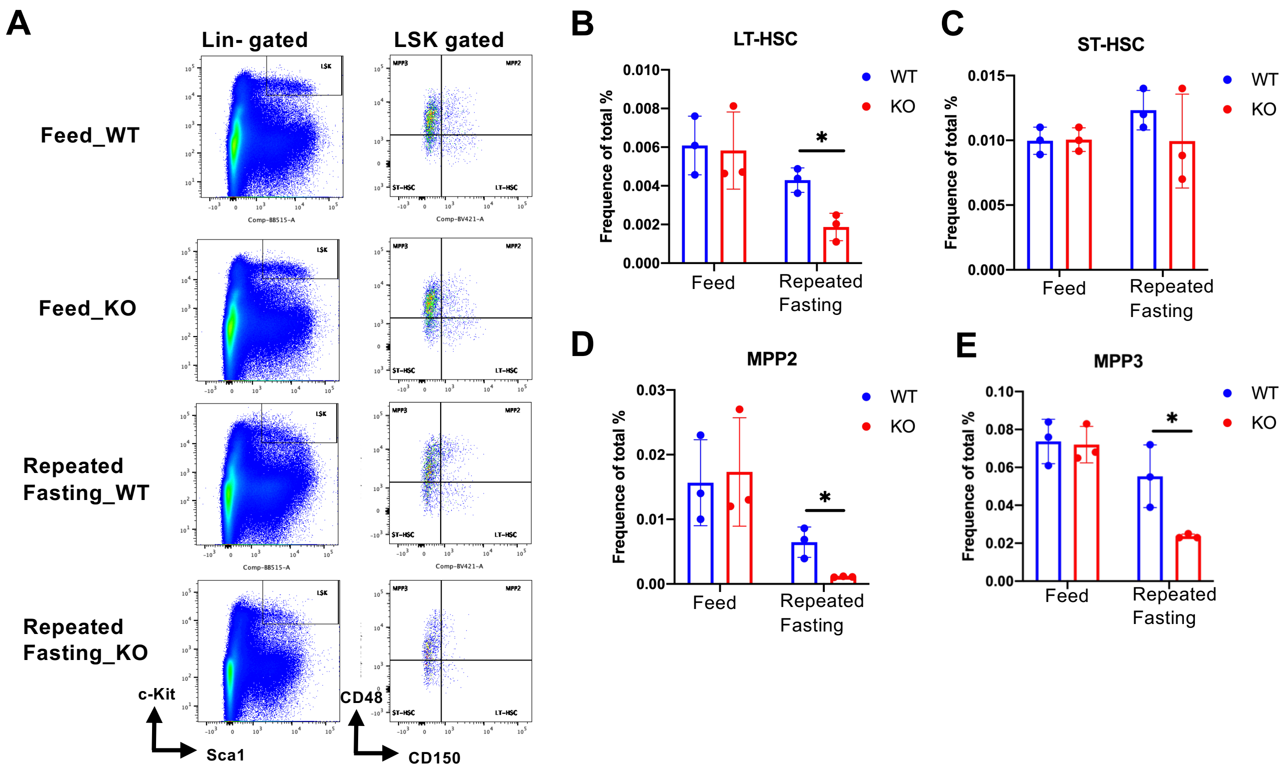

**Supplementary Table 1**

Mouse primers sequence information for quantitative PCR is listed as below. 5’→3’

| | | |
|---|---|---|
| *Hmgcs2* | Forward | GAAGAGAGCGATGCAGGAAAC |
| | Reverse | GTCCACATATTGGGCTGGAAA |
| *Acly* | Forward | CCCCAAGATTCAGTCCCAAGT |
| | Reverse | GCCTTGGTATGTCGGCTGAA |
| *Fasn* | Forward | CCCACTCTGGTTCATCTGCTC |
| | Reverse | CTTTCACTCCCAACGGCTTC |
| *Acaca* | Forward | CGCAAATGTGGAGTTGATTCTTG |
| | Reverse | AGTTCTGGGAGTTTCGGGTTC |
| *Acss2* | Forward | GTTGCTGCCTGCTCATCACTAC |
| | Reverse | CTTCTCTCGGCACTTCTCCAA |
| *Acat1* | Forward | CTATTTCCACTCCATGCACCAC |
| | Reverse | CCTGCCACCATCACATCCT |
| *Scd1* | Forward | GGAAAGTGAGGCGAGCAAC |
| | Reverse | GTGGTGGTGGTCGTGTAAGAA |
| *Pdk1* | Forward | TGGAAGTCAGTTTGGAATCAGG |
| | Reverse | AGAAGATTGTCGGGGAGGAG |
| *Slc25a1* | Forward | TCCGGGAAGCCCAAACT |
| | Reverse | ACGTATTCGGTCGGGAAGG |
| *Hmgcr* | Forward | CCTGGGAAGTTATTGTGGGAAC |
| | Reverse | GGATGATGATGTCGCTGCTC |
| *Oxct1* | Forward | GGGTGTGTCTGCTACTTCTCTGTC |
| | Reverse | AAAACCACCAACCAGCAAGG |
| *Gapdh* | Forward | CTTTGTCAAGCTCATTTCCTGG |
| | Reverse | TCTTGCTCAGTGTCCTTGC |

#### Source Data

This Western Blot result is relevant to Fig.4G

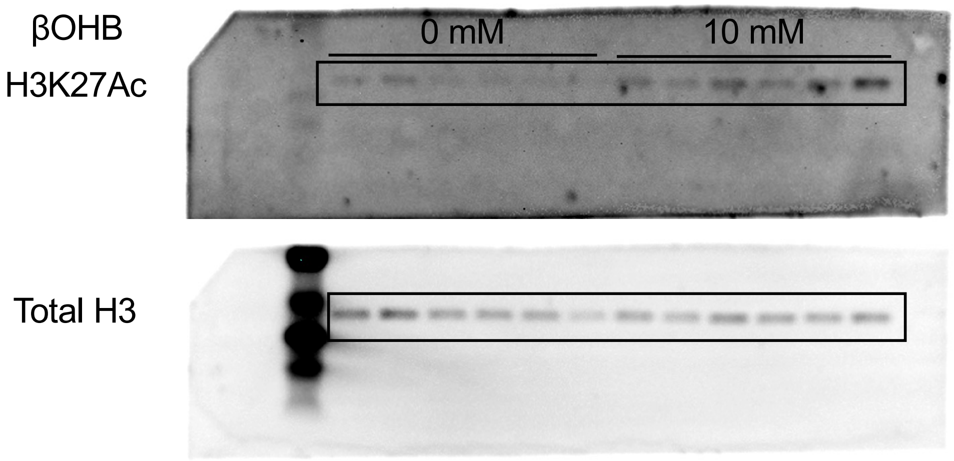
